# Flexible males, proactive females: personality tests show increased boldness/exploration in colonists damping with time in males but not in females

**DOI:** 10.1101/2024.05.22.595322

**Authors:** Andrey V. Tchabovsky, Elena N. Surkova, Ludmila E. Savinetskaya, Ivan S. Khropov

**Affiliations:** Laboratory for Population Ecology, Severtsov Institute of Ecology and Evolution, Russian Academy of Sciences, 33 Leninskii pr., Moscow, Russia

**Author notes:** Andrey V. Tchabovsky:., Elena N. Surkova:., Ludmila E. Savinetskaya:., Ivan S. Khropov. Corresponding author: A.V. Tchabovsky. Correspondence: Laboratory for Population Ecology, A.N. Severtsov Institute of Ecology and Evolution, Russian Academy of Sciences, 33 Leninskii pr., Moscow 119071, Russia.

**Keywords:** colonization, behavioural traits, phenotypic plasticity, gerbils, landscape change, range expansion

## Abstract

Individuals colonizing new areas at expanding ranges encounter challenging and unfamiliar environments, suggesting that colonists should differ in behavioural traits from the residents of the source populations. The colonizer syndrome is supposed to be associated with boldness, exploration, activity, and low sociability. We assessed spatial and temporal variation in the colonizer syndrome in the expanding population of midday gerbils (*Meriones meridianus*). Male first colonists tended to be faster and bolder than residents, although the difference was not significant. Female first colonists were bolder, faster and more explorative than residents. These findings support boldness/exploration syndrome as a typical colonizer trait, which appears to be restricted to females in midday gerbils. Males and females also differed in the behavioural dynamics post-colony establishment. In males, boldness/exploration/sociability peaked in newly founded colonies, then sharply decreased in subsequent generations following decreasing environmental uncertainty in aging colonies. In females, increased boldness/exploration did not lower with time post-colonization, i.e. female colonists retained bold/explorative phenotype in subsequent generations despite facing a less challenging environment. Thus, female colonists, unlike males, carry a specialized behavioural colonizer phenotype corresponding to a proactive behavioural coping strategy. We link sex differences in behavioural traits of colonists to the sex-specific life-history strategies.

## Introduction

The propensity to disperse and colonize new areas is believed to be associated with a set of individual traits, genetically and(or) developmentally determined, that distinguish dispersers and colonists from the residents of the source population (Myers, Krebs 1971; Bekoff 1977; Clobert et al. 2001, 2009; Krackow 2003; Ronce 2007; Matthysen 2012; Carere, Gherardi 2013; Chuang, Peterson 2016; Duckworth et al. 2018). If these traits are correlated, they form a specific dispersal syndrome (Cote et al. 2010a; Ronce, Clobert 2012; Sih et al. 2012). The propensity to disperse and the ability to transfer to and settle in a new location were shown to correlate with boldness (risk-taking), exploration (novelty seeking), activity, low sociability, and high aggressiveness (Dingemanse et al., 2003; Duckworth, Badyaev 2007; Duckworth, Kruuk 2009; Cote et al. 2010b; Debeffe et al. 2014; Liebl, Martin 2012, 2014; Duckworth et al. 2018) – personality traits associated with the proactive strategy of copying with environmental challenges (Koolhaas et al. 1999; Carere et al. 2010; Coppens et al. 2010; Baugh et al. 2017). For example, in Great tits (*Parus major*), natal dispersal distances correlated positively with the exploratory behaviour of individuals and their parents, implying that dispersal and novelty-seeking are linked and heritable (Dingemanse et al. 2003). Furthermore, the levels of exploratory and risky behaviours in Great tits appeared to be underpinned by a polymorphism in D4 dopamine receptor gene Drd4 (Fidler et al. 2007 but see Korsten et al. 2010) associated with novelty-seeking and migration distances in humans that colonized the South America (Tovo-Rodrigues et al. 2010). Dark-eyed juncos (*Junco hyemalis*) colonizing a novel urban environment in Southern California were bolder and more explorative than birds from the source population (Atwell et al. 2012). In western bluebirds, *Sialia mexicana*, aggressive males dispersed from the core populations to the front of invasion in the northwestern United States where displaced local less aggressive mountain bluebirds, *Sialia currucoides*, promoting further range expansion (Duckworth, Badyaev 2007). Moreover, the aggression of western bluebirds-colonists decreased rapidly across consecutive generations due to the selection on heritable low-flexible aggressive phenotypes after colony establishment (Duckworth, Kruuk 2009). The negative relationship between sociability and the propensity to disperse was shown for females of the yellow-bellied marmot *Marmota flaviventris* (Blumstein et al. 2009) and the greyLsided vole *Myodes rufocanus* (Ims 1990) as well as in the invasive mosquitofish, *Gambusia affinis,* (Cote et al. 2010b), supporting the social cohesion hypothesis by M. Bekoff (1977). The existence of consistent specific behavioural traits in colonists indicates that they are not just a random sample from the source population but spatially sorted phenotypes with bolder, more explorative or more aggressive individuals found at the edges of expanding ranges where they face novel and uncertain ecological and social conditions (Duckworth, Badyaev 2007; Duckworth, Kruuk 2009; Clobert et al., 2009; Chuang, Peterson 2016).

Nevertheless, dispersal-related behavioural traits are not always revealed or follow the predicted direction. For example, in the round goby *Neogobius melanostomus*, fish from the recently established populations were less bold than fish from the older population (Galli et al. 2023). In gambusia (*Gambusia affinis*), the syndrome of correlated individually consistent traits supposed to be associated with the propensity and ability to disperse (general activity, exploration, and boldness) was not related to the dispersal distance (Cote et al. 2010b). Real-time observations of invasive bank voles (*Myodes glareolus*) colonizing new areas during the ongoing range expansion in Ireland showed that males, the dispersing sex, were less bold at the expansion edge (Eccard et al. 2023).

The inconsistencies in the results may be attributed to the lack of field studies of ongoing colonization processes observed in real time: very few studies investigated not only spatial variation in behavioural traits across expanding ranges but also the temporal dynamics of behavioural profiles post-colonization (see for example, Duckworth, Kruuk 2009; Eccard et al. 2023). In the lack of direct observations in real time, the evidence for behavioural sorting during colonization remains equivocal. Additionally, it is unclear which behavioural types are favourable in different spatial zones of expanding ranges and at different stages of colonization (Eccard et al. 2023). Colonization begins with the arrival of the first colonists settling in the vacant area, settlers, followed by the flow of joiners, the immigrants from the core population (Clobert et al. 2009; Cote et al. 2010a). Settlers, immigrants, and their recruited offspring face different ecological and social challenges, as environmental novelty and uncertainty decrease with time after first arrivals with increasing habitat familiarization, social cues, and social landscape connectivity (Wey et al. 2015; Buxton et al. 2020). This implies that colonists’ behaviour may not only differ consistently from that of the residents of the source population due to spatial sorting but also may change with the colony’s age. This change may occur due to the adaptation to local ecological and social conditions in colonies in the long-term perspective (such as in the western sialia – Duckworth, Kruuk 2009; Duckworth et al. 2015), as well as in the short-term perspective because of behavioural flexibility. Real-time observations can provide an opportunity to follow these changes, which otherwise can be missed (Galli et al. 2023; Tchabovsky et al. 2023a).

Recent dynamics of rodent populations in the rangelands of Kalmykia (southern Russia) driven by human-induced landscape transformations provide an example of rapid species range shifts following cycles of desertification-steppification (Neronov et al. 1997; Shilova et al. 2000; Hölzel et al. 2002; Rogovin 2007; Dubinin et al. 2011; Smelansky, Tishkov 2012; Tchabovsky et al. 2023b). In particular, in response to the expansion of the tall-grass steppe in the 1990s, the population of the desert-dwelling midday gerbil, *Meriones meridianus*, first abruptly declined and then collapsed, and by mid-2010s, its range contracted to the east (Tchabovsky et al. 2016, 2023b). The new cycle of desertification launched the expansion of the midday gerbil population in the opposite direction – to the west in 2019, where they colonize re-emerging desert habitats, providing a rare opportunity to observe the colonization process in real time.

In this study, we aimed to assess spatial and temporal variation in behavioural traits of midday gerbils in the expanding population using multi-assay personality tests previously designed to measure boldness, exploration, sociability, and docility in gerbils (Tchabovsky et al. 2024). We hypothesized that the first colonists would differ from the residents of the source population in higher boldness and exploration and lower docility. We did not have any specific predictions on the sociability of colonists because previous results and the theory on the role of sociability in colonization success are most controversial (Duckworth et al. 2018). Further, we suggested that if the behavioural traits of colonists would quickly change with the colonies’ age approaching the baseline characteristics of the residents of the source populations, this would indicate a flexible behavioural response to decreasing uncertainty of social and ecological conditions. If, on the contrary, behavioural profiles of colonists would remain specific and consistent over generations, this would indicate spatial sorting of specialized colonizer phenotypes.

## Material and Methods

### Model species

*Meriones meridianus* is a small (50-60 g of adult body mass), short-lived (ca eight months of life expectancy) nocturnal-diurnal mainly granivorous psammophilous gerbil inhabiting deserts and semi-deserts of Central Asia, Northern China, and Southern Russia (Wang et al. 2013; Nanova et al. 2014; Tchabovsky et al. 2019a; Yang et al. 2020). Midday gerbils live solitarily or in loose aggregations: males and females are not territorial, do not form pair bonds, are socially promiscuous, interact rarely, display little agonistic or amicable behaviour, and, on the whole, are socially indifferent (Gol’tsman et al. 1994; Tchabovsky et al. 2004; Shilova, Tchabovsky 2009; Tchabovsky et al. 2019b). Breeding season in Kalmykia typically lasts from early April to mid-September. In mid-April–mid-May, when the tests were performed, the population consisted mainly of overwintered adults and some young (excluded from the analysis due to small sample sizes).

### Study area

The study area is located in the semi-desert rangelands of southeastern Kalmykia (southern Russia) and the Astrakhanskii region adjacent to the east. Kalmykia represents the very northwestern edge of the species range, where the population dynamics of gerbils follow the cycles of desertification-steppification (Shilova et al. 2000; Tchabovsky et al. 2016, 2019a, 2023b). We subdivided the entire area into two zones: the Western zone, at the edge of the species range, and the Eastern zone, extending towards the range core. The Western zone spans 50 km from the range border in the west to the east and features more or less isolated sandy patches suitable for gerbils that have been monitored annually twice a year since 1994 (in mid-April–mid-May and mid-September–mid-October). In 2017, the local population had collapsed, and gerbils disappeared from the entire Western zone. We continued monitoring the abandoned Western zone twice a year to be prepared for the gerbils to come back following the new cycle of desertification (Tchabovsky et al. 2023b). The Eastern zone, where gerbils have persisted constantly since the 1960s (Varshavsky et al. 1991, Tchabovsky et al. 2023b), extends further to the east and northeast, towards the range core, for another 100 km and has been monitored since 2017. The invasion of gerbils into the Western zone began in the autumn of 2019, with the first few colonists appearing in the two sites at its eastern borders. Since then, new colonies have emerged in the Western zone every year, discovered by the regular inspections of all monitored locations during spring and autumn sessions (Tchabovsky et al. 2023b).

### Trapping procedure

In this study, we used data collected in 2022-2023 in spring (mid-April – mid-May). We trapped gerbils with modified non-commercial Shchipanov’s wired live traps (Shchipanov 1987) baited with sunflower seeds and placed at the entrances of active burrows at the beginning of evening activity (about 19:00). We set traps for 3-5 hrs and checked them once an hour. Captured gerbils were carried in traps to a mobile field laboratory located 50-200 meters away, where tests and other manipulations were performed.

### Experimental procedure

The entire procedure designed to assess personality traits in gerbils (Tchabovsky et al. 2024) included four consecutive tests (Table 1). The first test evaluated a single measurement – the activity of a gerbil in a handling bag. The second, third, and fourth tests were composite, included multiple measurements, and were carried out in suits of consecutive assays within the same experimental arena.

**Table 1.**
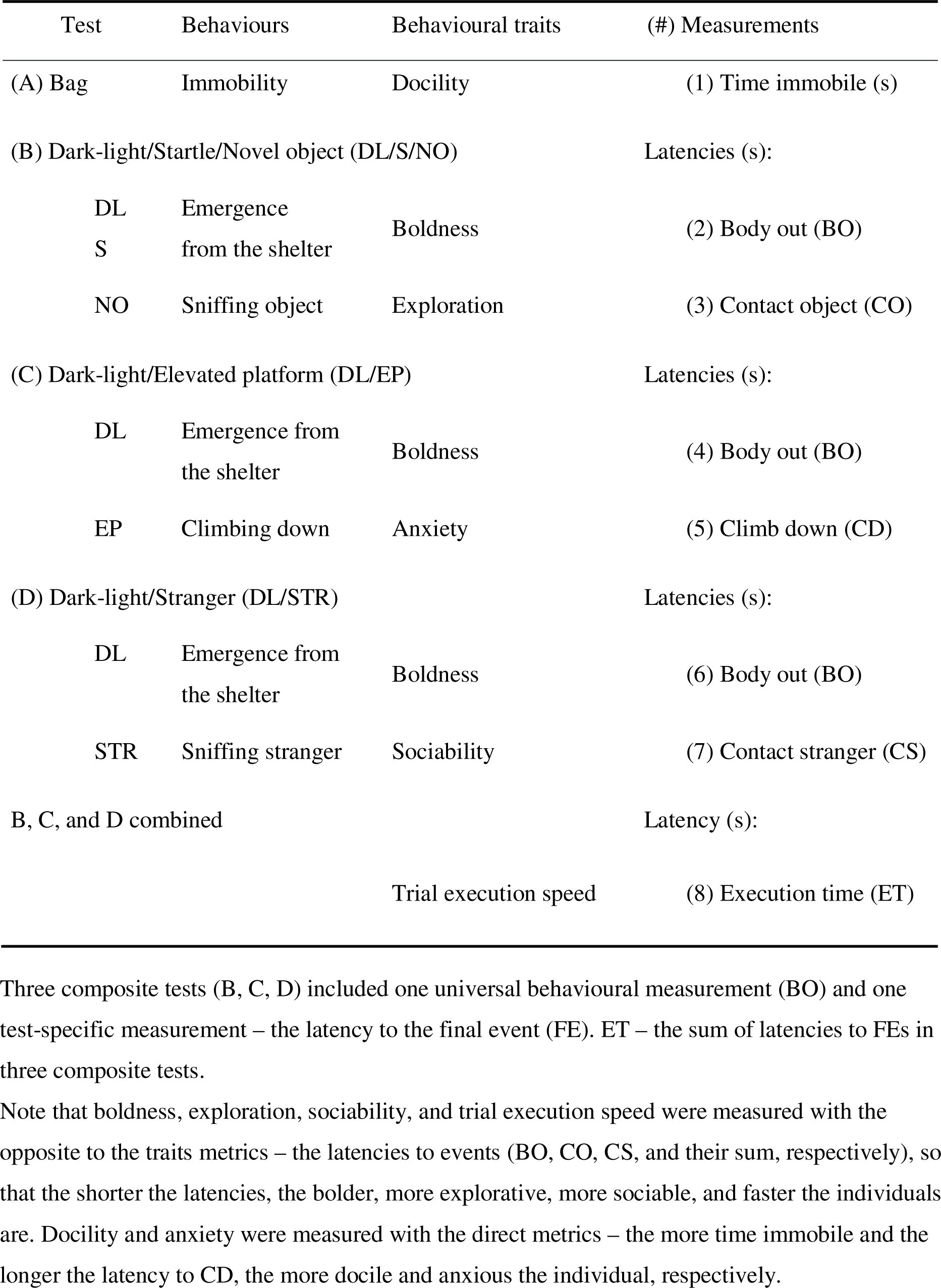
The sequence of tests, recorded behaviours, associated traits, and measurements.

First, to assess docility (response to restraint) we placed a small cotton handling bag (35 x 25 cm) with a gerbil inside in a plastic bowl and measured the time (in seconds) the gerbil spent immobile during 1 min. Then the animal was released into an opaque metal box (25 x 7 x 7 cm) placed in the illuminated plastic arena (78 x 56 x 43 cm) with a floor covered with sand. After 1 min of acclimation we opened the door of the box distantly to start the dark-light (DL) Startle/Novel object test. When the gerbil showed its head from the box, we dropped with noise a metal cylinder (5 x 4 cm) at the opposite side of the arena and recorded the latency from the start of the experiment until the gerbil emerged from the box with all four limbs (“body out” – BO) as a measure of boldness and then the time it took the gerbil to approach and sniff the dropped cylinder (“contact novel object” – CO) as a measure of exploration.

At the next stage, we returned the gerbil to the box and placed it on the elevated (16 cm) Plexiglas transparent platform (50 x 8 cm) with no walls within the same illuminated arena. After 1 min of habituation, we opened the door and recorded the latency to “body out” event and then the time it took the gerbil to climb down (CD) as a measure of anxiety (fear-related behaviour). At the final stage we removed the platform, returned the gerbil to the box and placed a wire-mesh cage (15 x 6.5 x 6.5 cm) with an unfamiliar adult male at the side of arena, opposite the box with a focal gerbil. After 1 min of habituation, we opened the door and measured the latency to the “body out” event and then the time it took the gerbil to approach and contact (sniff) the stranger through the wire mesh (CS) to assess sociability. If an animal did not show full body from the box or did not climb down, approached the object or stranger during 5 min of the test, the latencies were set to 300 s. At the end of the trial, the setup and equipment were cleaned with 70% Ethanol, and the floor of the arena was covered with fresh sand before introducing the next animal. In total we performed 180 tests (87 for 71 males and 93 for 73 females). More details of the experimental procedure can be found elsewhere (Tchabovsky et al. 2024).

### Data analysis

#### Classification of colonies

We fixed a colonizing event when gerbils first appeared in spring or the following autumn in the location that had remained vacant since 2017. Adult gerbils in newly founded colonies represented the first settlers (the true colonists). The colonies of ages 1, 2, and 3, i.e. the same colonies in subsequent years after the founding event, normally included no settlers and consisted of increasing numbers of locally-born individuals and, probably, new immigrants. In addition, with time post-colony foundation, the development of the social landscape and burrow network should make the environment more predictable and safer (Surkova et al. 2024). Therefore, new colonies differed intrinsically from the source population and the colonies of age 1+ in the composition, which consisted of first settlers, and by higher environmental uncertainty and novelty. Thus, to assess spatio-temporal variation in behavioural traits during colonization, we compared behavioural measurements between gerbils residing in the source population, new colonies, and the colonies of ages 1 and 2-3. We pooled data for colonies of ages 2 and 3 because the sample for colonies of age 3 was small.

#### Statistical analysis

First, we tested for the differences in eight primary behavioural measurements (Table 1) between residents of the source population and first colonists, i.e. gerbils from the newly established colonies. If an animal did not appear from the box at all (did not show its head) during any stage of the trial in the arena, we excluded this stage from the analysis of respective variables. In addition, we excluded measurements for the “climb down” event if a gerbil just fell from the platform. Thus, sample sizes may vary between primary variables. For the trial execution time we excluded one gerbil that did not appear (did not show head) at all four stages, but included others who appeared at least at one stage.

Then, to analyse the temporal dynamics of behavioural traits during colonization, we used composite variables obtained with principal component analysis (PCA) from a set of original behavioural measurements, a common and valid tool to reduce the dimensionality of multicollinear datasets in behavioural research (Budaev 2010; Mazzamuto et al. 2019). PCA was conducted separately for males and females with the complete data set with no missing values for any of the primary variables (64 tests for 54 males and 76 for 56 females) (Table 1; trial execution time was not included in PCA as a combination of three variables – the latencies to final events). PCs were interpreted as composite behavioural responses according to the loadings of primary variables (> 0.6 in absolute value as recommended for small sample sizes – Budaev 2010). Each animal was assigned scores from each of the retained PCs. PC scores were used as response variables to compare behavioural profiles between gerbils from the source population, new colonies, and the colonies of ages 1 and 2-3.

Statistical analysis was performed in R 4.2.3 (R Development Core Team, 2023). We assessed the variation of primary behavioural measurements as well as composite variables derived with PCA between the source population and colonies in adult males and females by constructing separate linear mixed-effect models (LMMs) with a restricted maximum likelihood (REML) method implemented in the *lme* function in the R package *nlme* (Pinheiro et al. 2021). Behavioural measurements and PCs were fitted to the models as response variables, while the residence category of gerbils (gerbils from the source population, new colonies, and the colonies of ages 1 and 2-3) was included as a predictor. We mainly aimed to assess the differences in behaviour between newly emerged colonies and source population as well as the changes in behaviour with the colony age post foundation. Thus, we considered the new colonies as a reference group. Given we sampled some individuals two times (8-25% depending on the test type), we fitted the animal identity to the models as a random term. All measurements prior to analyses were ln(*x*+1) transformed to improve normality.

## Results

### New colonies versus source population

Time spent immobile in the bag test did not vary between the new colonies and the source population, either in males or females (Table 2). In males, the latencies to events tended to be shorter for colonists compared to the source population (i.e. colonists tended to be faster) at all stages of the composite test, except the latency to climb down from the elevated platform (Fig. 1). However, the difference approached significance only for the “contact stranger” event (Table 2). Females from new colonies compared to their counterparts from the source population were faster at all stages of the composite test and, unlike males, exhibited significant differences in many measurements. They were bolder as indicated by the significantly shorter latencies to show full body at the dark-light stages of the startle and the stranger tests as well as in the elevated platform test (the difference is close to significance). They also tended to be faster in exploring the novel object and were significantly faster in climbing down from the elevated platform and executing the entire trial (Table 2, Fig. 1). Random effects of animal ID were insignificant in all models except for immobility in males (L-ratio = 17.3, p < 0.0001).

**Figure 1.**
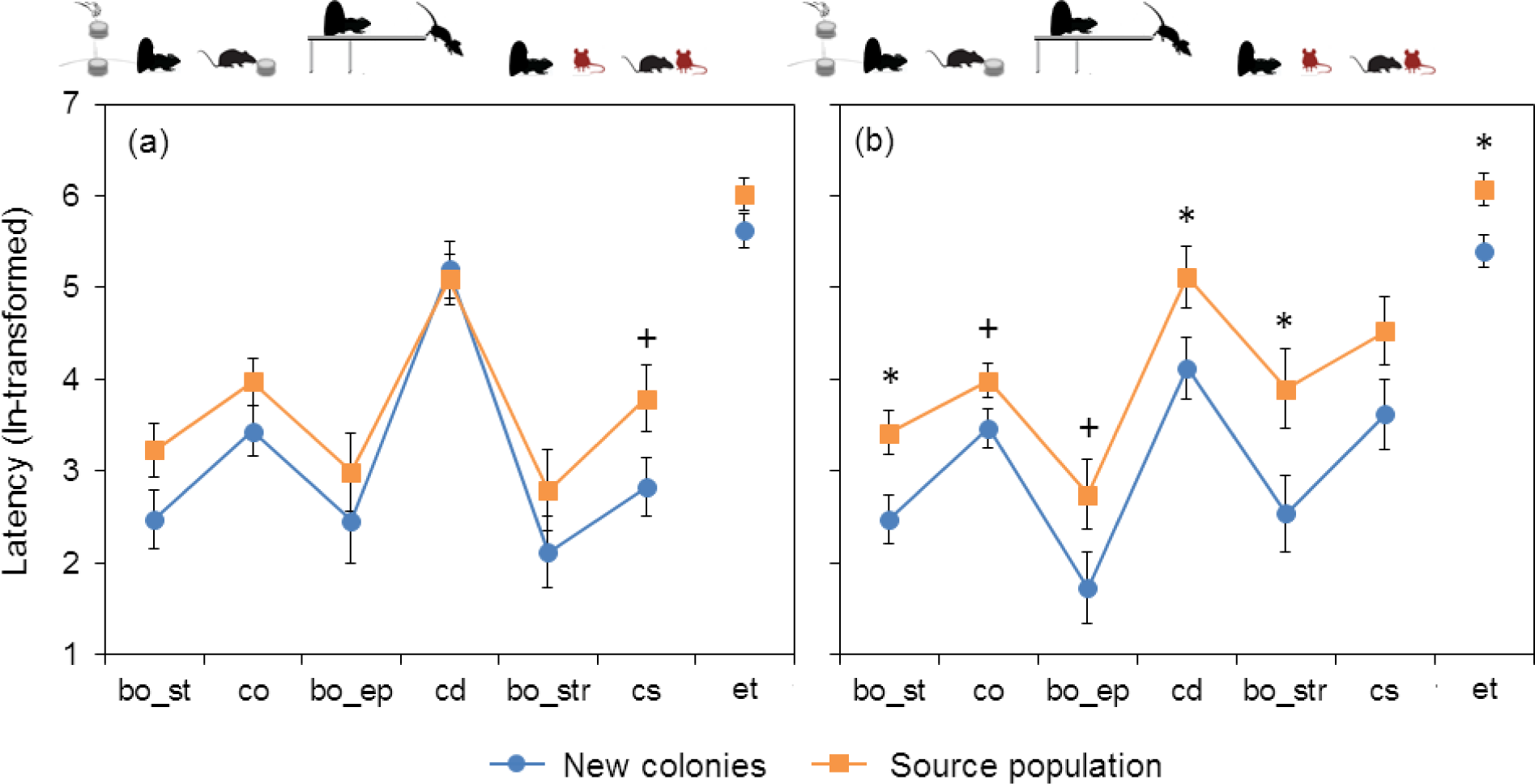
Mean ± SE latencies (ln-transformed) to consecutive events in the composite tests (see Table 1) for males (a) and females (b) from the new colonies and the source population. Asterisks and plus signs indicate significant and close-to-significance differences, respectively, as revealed by LMMs (see Table 2). bo – body out, co – contact object, cd – climb down, cs – contact stranger. et – trial execution time. st – startle test, ep – elevated platform test, str – stranger test.

**Table 2.** Coefficients of the linear mixed-effects models (*B*□±□SE) assessing the effect of the residence category of gerbils (the source population or new colonies) on behavioural measurements in adult males and adult females. The numbers of tests and tested individuals are shown in brackets.

| Fixed effects | Adult males |  |  | Adult females |  |  |
| --- | --- | --- | --- | --- | --- | --- |
| | $B \pm SE$ | $t$ | $p$ | $B \pm SE$ | $t$ | $p$ |
| IMMOB (s) | $0.03 \pm 0.09$ (39/31) | 0.3 | 0.7 | $-0.04 \pm 0.12$ (39/32) | -0.3 | 0.7 |
| BO_ST (s) | $0.76 \pm 0.43$ (37/30) | 1.8 | 0.09 | **$0.94 \pm 0.36$ (39/32)** | **2.6** | **0.01** |
| CO (s) | $0.55 \pm 0.38$ (37/30) | 1.4 | 0.2 | <u>$0.51 \pm 0.27$ (39/32)</u> | <u>1.9</u> | <u>0.07</u> |
| BO_EP (s) | $0.54 \pm 0.63$ (35/28) | 0.8 | 0.4 | <u>$1.02 \pm 0.55$ (37/30)</u> | <u>1.9</u> | <u>0.07</u> |
| CD (s) | $-0.11 \pm 0.41$ (31/25) | -0.3 | 0.8 | **$0.99 \pm 0.47$ (36/29)** | **2.1** | **0.046** |
| BO_STR (s) | $0.67 \pm 0.58$ (29/24) | 1.1 | 0.3 | **$1.36 \pm 0.59$ (33/26)** | **2.3** | **0.03** |
| CS (s) | <u>$0.95 \pm 0.48$ (29/24)</u> | <u>2.0</u> | <u>0.058</u> | $0.91 \pm 0.52$ (33/26) | 1.7 | 0.1 |
| ET (s) | <u>$0.40 \pm 0.26$ (38/31)</u> | 1.5 | 0.1 | **$0.67 \pm 0.25$ (39/32)** | **2.7** | **0.01** |

IMMOB – time immobile in the bag, BO – body out, CO – contact object, CD – climb down, CS – contact stranger, ET – trial execution time, ST – startle test, EP – elevated platform test, STR – stranger test.
Note that boldness, exploration, sociability, and trial execution speed were measured with the opposite to the traits metrics – the latencies to events (BO, CO, CS, and their sum), so that the shorter the latencies, the bolder, more explorative, more sociable, and faster the individuals are.
Docility and anxiety were measured with the direct metrics – the more time immobile and the longer the latency to CD, the more docile and anxious the individual, respectively.
New colonies were the reference group, s – source population. Significant and close to significance effects are marked with bold and underlined fonts, respectively. Animal ID was added as a random term to all models.

### Temporal dynamics of behavioural traits post-colonization

PCA of primary behavioural measurements conducted separately for males and females produced three PCs with eigenvalues ≥ 1.0 for each sex, which in total explained 71% and 74% of the variance, respectively (Table 3). In males, PC1 combined short latencies to “body out” in the startle and stranger tests as well as to contact the novel object and the stranger and was labelled “Boldness/Exploration/Sociability”. PC2 was mainly associated with the short latency to climb down from the platform indicating low anxiety and was termed “Confidence”. PC3 was mainly associated with low activity in the bag, i.e. reflected docility. In females, PC1 combined short latencies to show full body in dark-light emergence assays in all three contexts and the short latency to contact object and was labelled “Boldness/Exploration”. PC2 was mainly associated with the short latency to contact a stranger and additionally to show body in the same test, i.e. reflected sociability, while PC3 was correlated with the short latency to climb down, i.e. indicated low anxiety and was termed “Confidence”

**Table 3.** Results of the PCA (PCA loadings) of primary behavioural measurements.

| Measurements | Adult males |  |  | Adult females |  |  |
| --- | --- | --- | --- | --- | --- | --- |
|  | PC1 | PC2 | PC3 | PC1 | PC2 | PC3 |
| IMMOB | 0.02 | -0.45 | **0.79** | -0.44 | 0.18 | 0.50 |
| BO_ST | **-0.79** | 0.40 | 0.24 | **-0.81** | 0.28 | 0.13 |
| CO | <u>-0.59</u> | 0.54 | 0.16 | **-0.81** | 0.27 | 0.22 |
| BO_EP | -0.25 | -0.51 | -0.50 | **-0.64** | <u>0.43</u> | -0.27 |
| CD | -0.17 | **-0.69** | 0.20 | -0.45 | 0.07 | **-0.77** |
| BO_STR | **-0.84** | -0.16 | -0.06 | **-0.71** | <u>-0.59</u> | -0.02 |
| CS | **-0.83** | -0.32 | -0.15 | <u>-0.58</u> | **-0.72** | 0.04 |
| Eigenvalue | 2.5 | 1.5 | 1.0 | 3.0 | 1.3 | 1.0 |
| % Total variance | 35.0 | 21.7 | 14.7 | 42.3 | 17.9 | 14.0 |

| PC label | Boldness/<br>Exploration/<br>Sociability | Confidence | Docility | Boldness/<br>Exploration | Sociability | Confidence |
| --- | --- | --- | --- | --- | --- | --- |

PCA loadings > 0.6 and close to 0.6 in absolute value are in bold and underlined fonts, respectively. Pairwise correlations between primary variables did not exceed 0.8, and all of them were included in the analysis.
IMMOB – time immobile in the bag, BO – body out, CO – contact object, CD – climb down, CS – contact stranger. ST – startle test, EP – elevated platform test, STR – stranger test. Note that boldness, exploration, and sociability were measured with the opposite to the traits metrics – the latencies to events (BO, CO, and CS, respectively), so that the shorter the latencies, the bolder, more explorative, and more sociable the individuals are. Docility and anxiety were measured with the direct metrics – the more time immobile and the longer the latency to CD, the more docile and anxious the individual, respectively. Therefore, the negative (positive) loadings of the relevant primary variables (BO, CO, and CS) indicate positive (negative) correlations with respectively boldness, exploration, and sociability. The signs of loadings for immobility and CD indicate the direction of correlation with docility and anxiety, respectively.

In males, PC1 scores (Boldness/Exploration/Sociability) varied with the colony age, being significantly lower in colonies of age 2-3 than in new colonies, where they were the highest (Table 4, Fig. 2). Although PC1 scores were also higher in new colonies as compared to the source population, the difference was insignificant. PC2 (Confidence) and PC3 (Docility) did not vary between the new colonies and source population or between new and older colonies. In females, PC1 scores (Boldness/Exploration) were significantly higher in new colonies than in the source population but did not significantly change with the colony age. PC2 (Sociability) did not vary across residence categories of females, whereas PC3 (Confidence) was significantly higher in new colonies than in older colonies and close to significance higher than in the source population. Random effects of animal ID were insignificant in all models.

**Figure 2.**
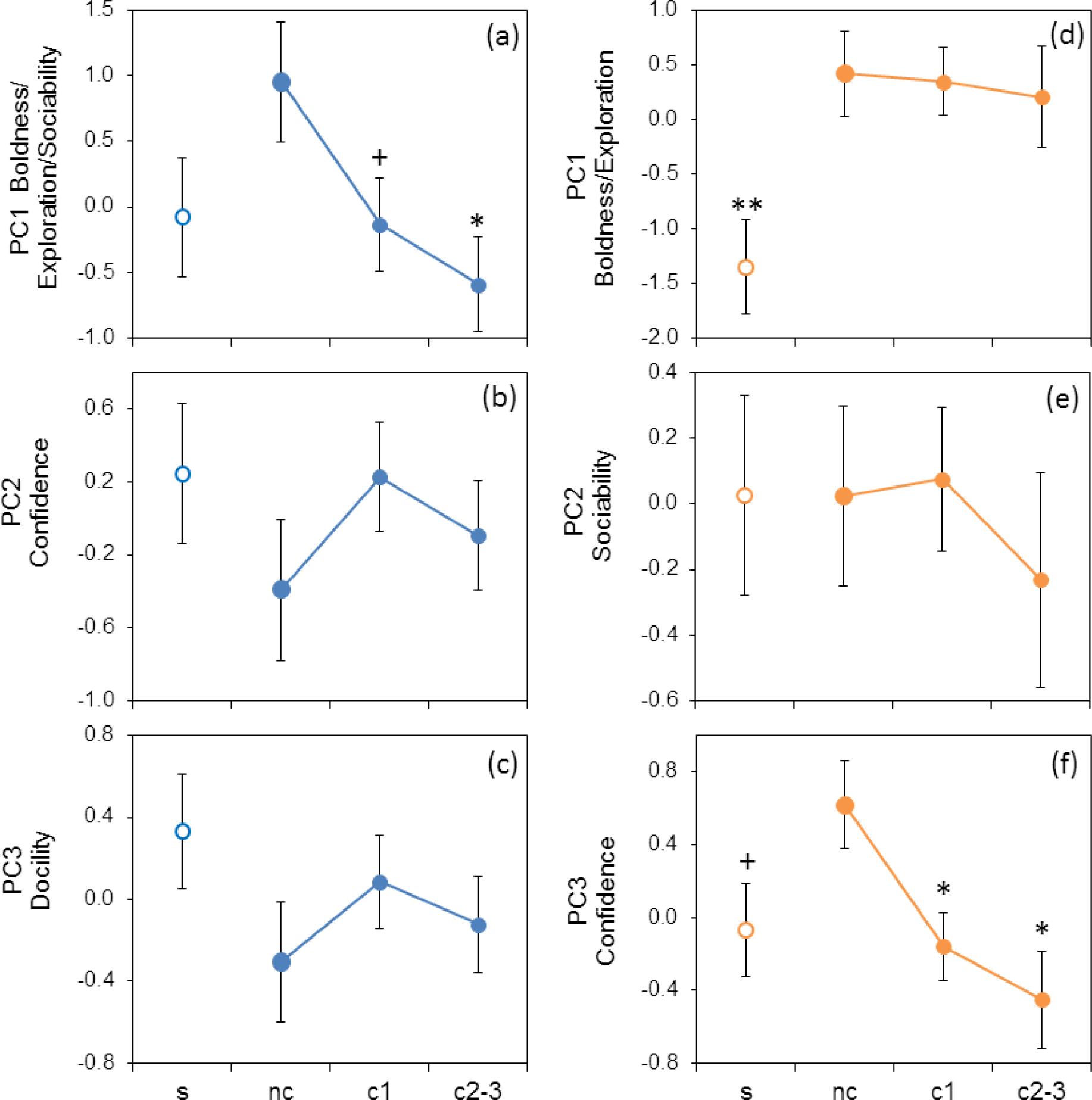
Variation in behavioural traits between the source population (s, open markers), newly founded colonies (nc), and colonies of ages 1 (c1) and 2-3 (c2-3) for adult males (a-c) and adult females (d-f). Mean ± SE for PCs scores (see Table 3). Enlarged markers indicate the reference group (nc) in LMM. Asterisks and plus signs show significant and close to significance contrasts with the reference group, respectively.

**Table 4.** Coefficients of the linear mixed-effects models (*B*□±□SE) assessing the effect of the residence category of gerbils (the source population, new colonies, colonies of age 1, and colonies of age 2-3) on the composite behavioural traits derived from PCA (see Table 3) in adult males and adult females. The numbers of tests and tested individuals are shown in brackets.

| Adult males (64/54) | Adult females (70/56) |
| --- | --- |

| Fixed effects | $B \pm SE$ | $t$ | $p$ | $B \pm SE$ | $t$ | $p$ |
| --- | --- | --- | --- | --- | --- | --- |
|  | PC1 - Boldness/Exploration/Sociability |  |  | PC1 - Boldness/Exploration |  |  |
| <u>c1</u> | <u>-1.09 ± 0.58</u> | <u>-1.86</u> | <u>0.068</u> | -0.07 ± 0.50 | -0.15 | 0.9 |
| c2-3 | **-1.54 ± 0.59** | **-2.63** | **0.01** | -0.21 ± 0.61 | -0.35 | 0.7 |
| s | -1.03 ± 0.65 | -1.59 | 0.1 | **-1.77 ± 0.58** | **-3.04** | **0.004** |
|  | PC2 - Confidence |  |  | PC2 - Sociability |  |  |
| c1 | 0.62 ± 0.49 | 1.26 | 0.2 | 0.05 ± 0.35 | 0.15 | 0.9 |
| c2-3 | 0.30 ± 0.49 | 0.60 | 0.5 | -0.26 ± 0.43 | -0.60 | 0.6 |
| s | 0.64 ± 0.55 | 1.17 | 0.2 | 0.00 ± 0.41 | 0.00 | 1.0 |
|  | PC3 - Docility |  |  | PC3 - Confidence |  |  |
| c1 | 0.39 ± 0.37 | 1.06 | 0.3 | **-0.78 ± 0.30** | **-2.56** | **0.025** |
| c2-3 | 0.18 ± 0.37 | 0.49 | 0.6 | **-1.07 ± 0.36** | **-2.96** | **0.01** |
| s | 0.64 ± 0.41 | 1.57 | 0.1 | <u>-0.68 ± 0.35</u> | <u>-1.93</u> | <u>0.059</u> |

New colonies were the reference group. c1 – colonies of age 1, c2-3, colonies of age 2-3, s – source population. Animal ID was added as a random term to all models. Significant and close to significant effects are marked with bold and underlined fonts, respectively.

## Discussion

In this study, using real-time observations of the first stages of the colonization process during range expansion in the midday gerbil, we found sex-specific patterns in behavioural variation between the residents of the source population and the first colonists, as well as in the short-term temporal dynamics of the behavioural traits post-colonization. To our knowledge, this is the first study of behavioural patterns of colonists in mammals during the first stages of colonization conducted in real time.

### Spatial-temporal dynamics of behavioural traits in expanding population

Although, as expected, male first colonists tended to be faster and bolder than residents of the source population at almost all stages of the behavioural tests, they did not show pronounced differences in primary measurements or composite behavioural traits (Tables 2 and 4, Figs 1 and 2). They differed from residents singly – being some faster in contacting a stranger, i.e. exhibited lower social shyness. In contrast to males, female first colonists differed consistently from the residents in being bolder and more confident in different contexts (both non-social and social), as indicated by the shorter latencies to emerge in dark-light assays in all three tests and the latency to climb down from the platform. They were also faster in inspecting a novel object and executing the entire trial (Table 2, Fig 1). These differences in primary measurements are supported by the analysis of composite variables: females-first colonists exhibited higher scores in boldness/exploration and confidence than residents of the source population (Table 4, Fig. 2). These results are consistent with our predictions and support boldness/exploration syndrome as a specific trait of dispersers and colonists universal for many – though not all – species (Dingemanse et al. 2003; Cote et al. 2010a; Debeffe et al. 2014; Liebl, Martin 2012, 2014; Chuang, Peterson 2016; Duckworth et al. 2018), which, in midday gerbils, turned out to be a female-specific characteristic. The opposite pattern was found in the other study of colonizing rodents, the bank voles, where colonist males were shyer than residents, whereas females showed no differences (Eccard et al. 2023). However, in that study, new colonies of ages 1-4 years were merged in a sample; therefore, short-term changes in behaviour post-colony establishment could be missed.

The lack of differences in sociability in females, combined with only a weak tendency for lower social shyness in male colonists, adds to the equivocal results on the role of individual sociability in colonization success (Duckworth et al. 2018; Tchabovsky et al. 2023a), which may vary between species due to species specificity of social behaviour. We previously have shown that sociability is a consistent personality trait in *M. meridianus* (Tchabovsky et al. 2019b, 2024), which, according to this study, seems unrelated to the colonization process. We attribute this to the overall low sociability as a species-specific trait of this socially indifferent gerbil (Gol’tsman et al. 1994; Tchabovsky et al. 2004; Shilova, Tchabovsky 2009). In particular, midday gerbils showed no preferences between familiar and unfamiliar conspecifics in the preference tests (Tchabovsky et al. 2019b), suggesting that their response to social novelty in colonies may be weak.

Males and females exhibited sex-specific patterns not only in the differences between the source population and the first colonists, strong in females and weak in males, but also in the temporal dynamics of behavioural traits post-colony establishment (Fig. 2). In males, boldness/exploration/sociability peaked in newly founded colonies, then sharply decreased in subsequent generations, becoming significantly lower 1-3 years post-foundation. This dynamic pattern follows the changes in environmental challenges and uncertainty, which should be the highest for the first settlers and decrease in subsequent generations of colonists with increasing habitat familiarization, social cues, and social landscape connectivity, as well as with the development of burrow network (Wey et al. 2015; Buxton et al. 2020; Surkova et al. 2024). The dynamic behavioural response of male colonists to changes in environmental uncertainty suggests that they do not carry specific colonizer phenotypes persisting across generations. Instead, it may reflect an adaptive phenotypic plasticity (Clobert et al. 2009; Dingemanse et al. 2010; Surkova et al. 2024), helping males explore, cope with, and settle in unfamiliar ecological and social environments at the first stage of colonization and then adjust behavioural responses to decreasing environmental uncertainty.

In female colonists, however, increased boldness/exploration did not lower with time post-colonization despite decreasing environmental uncertainty and the increasing number of locally born and recruited females (Fig. 2d). In other words, females have retained bold/explorative phenotype of the first settlers in subsequent generations despite facing a less challenging and uncertain environment. This implies that female colonists, unlike males, carry a rigid specialized behavioural colonizer phenotype corresponding to a proactive coping strategy (Koolhaas et al. 1999; Carere et al. 2010; Coppens et al. 2010; Baugh et al. 2017).

In contrast to boldness/exploration syndrome, confidence in the elevated platform test decreased with time in females after peaking in new colonies (Fig. 2f). Previously, we have shown that boldness in the dark-light startle test was positively correlated with confidence (as the opposite of anxiety) on the platform in *M. meridianus* in the laboratory, constituting a syndrome (Tchabovsky et al. 2024). However, in this study, boldness and confidence represent different behavioural axes in both sexes and differ in the pattern of post-colonization dynamics in females. Thus, confidence on the elevated platform appears to be a distinct form of risk-taking behaviour in colonists independent from boldness measured in other contexts. We do not have a clear explanation why later generations of female colonists become more cautious and timid in climbing down; whatever the reason, females exhibit plasticity in this behaviour post-colonization.

We found no variation in docility between the first colonists and residents or colonists of later generations in either males or females. This contradicts the hypothesis that population expansion selects for docile (non-aggressive) phenotypes that are more likely to colonize vacant habitats (Krebs et al. 1995). Previously, we have shown that docility (as a response to restrain) is not correlated with boldness in *M. meridianus* (Tchabovsky et al. 2024), consistent with other studies showing its independence from other personality traits (Petelle et al. 2013). Probably, docility, measured as a specific response to restrain, reflects a distinct behavioural trait irrelevant to colonization success.

### Plastic vs consistent colonist phenotypes of males and females

Dispersal and colonization processes include the same three stages: emigration, transfer, and settlement in a new location, but are distinct in required specific individual traits and costs, as well as in ecological and evolutionary consequences (Ims, Yoccoz 1997; Kerth, Petit 2005). Unlike dispersal within a population range, colonization is typically associated with long-distance movements beyond the genetic neighbourhood and(or) population geographic range (Bartoń et al. 2012; Jordano 2017) and often implies crossing the matrix in fragmented landscapes to settle in suitable but vacant from conspecifics habitats (Hanski 1994) – the gaps in the social landscape. Because in mammals, males and females differ in spatial behaviour and dispersal propensity, they may be exposed to differential selection pressures during the population expansion, producing sex-specific spatial sorting patterns in behavioural traits (Eccard et al. 2023). For example, in bank voles, timid and risk-averse males were found at the edge of the expanding range, whereas females were not behaviourally sorted (Eccard et al. 2023). The authors attributed these between-sex differences to sex-specific dispersal propensity: avoiding risks should be higher in males as dispersing sex and under strong selection during colonization of unfamiliar areas void of conspecifics.

In midday gerbils, we found a different pattern. First, males exhibited plastic behavioural profiles changing with time and were not behaviourally sorted between the core and edge populations, which contradicts the hypothesis on the spatial sorting of risk-averse males in an expanding population. Second, females did exhibit spatial sorting with less risk-averse and more explorative proactive phenotypes found in the edge population and retained in subsequent female generations of colonists. Eccard et al.’s idea that non-dispersing sex should not be selected for risk-aversion does not explain spatial sorting of typically philopatric females of midday gerbils, i.e. why female colonists are consistently bolder and more explorative than residents.

Like Eccard et al., we relate sex-specific patterns in plasticity and spatial-temporal dynamics of behavioural traits in an expanding population of gerbils to sex-specific life-history strategies and space use, but from another perspective. Sexual conflict theory assumes that contrary to the “risk-averse hypothesis”, males, unlike females, are selected for increased competitive abilities and risk-taking behaviour, which increases their reproductive success (Trivers 1972; Promislow 1992). Indeed, personality research shows that males are typically more explorative, bolder, and less risk-averse and experience higher predation risk than females (King et al. 2013, but see Harrison et al. 2022). In mammals, males are the dispersing sex, while dispersal is a risky process (Bonte et al. 2012). Together, this implies that males are likely to face more variable and less predictable environments, which may favour the flexibility of behaviour to respond adequately to immediate challenges.

In addition, in species with male-biased dispersal, females are often considered as the “colonizing sex” in contrast to males viewed as the “dispersing sex” (Kerth, Petit 2005; Gauffre et al. 2009). Colonizing females may differ in behaviour from females dispersing within the population because colonization requires specific individual traits and costs (Kerth, Petit 2005), in particular risk-taking and lower survival, especially in the fragmented landscape, as shown in the individual-based models (Bartoń et al. 2012; Markov, Ivanko 2022). Long-distance dispersal beyond genetic neighbourhood to occupy vacant habitats should be an especially difficult and costly challenge for normally phylopatric and kin-targeted mammalian females (Greenwood 1980; Lawson Handley, Perrin 2007; Hoogland 2013). Moreover, in the fragmented landscape of Kalmykian rangelands, crossing the matrix to settle in vacant from conspecifics sandy patches with unknown resources should be a special challenge for strongly philopatric site-attached females of the midday gerbil (Shilova, Tchabovsky 2009). Therefore, bold explorative colonizer phenotypes, stable over subsequent generations of colonist females and uncoupled from the decreasing environmental uncertainty, may be a female-specific individual trait favouring colonization.

Two sex-specific behavioural phenotypes of colonists – plastic in males and rigid proactive colonizer syndrome in females correspond to two alternative models explaining colonization and invasion success: behavioural flexibility and behavioural syndrome (Clobert et al. 2009; Cote et al. 2010a; Dingemanse et al., 2010; Chapple, Wong 2016; Chuang, Peterson 2016; Eccard et al. 2023). The former helps respond quickly to changing and unpredictable environmental conditions. The latter suggests that colonists are not just a random sample from the source population but behaviourally preadapted to face stressful and uncertain risky environments in colonized novel areas and, thus, spatially sorted. One essential inference from our study is that these two contrasting strategies may be sex-specific, with female colonists consistent while male colonists flexible in their behaviour.

Recently, we have shown similar sex-specific patterns in plasticity/consistency of stress regulation in the same colonizing population of gerbils: increased glucocorticoid levels in first colonists rapidly dampened with the colonies age in males but remained high across subsequent generations in females (Surkova et al. 2024). Therefore, in stress regulation, female colonists showed a reactive strategy associated with high activity of hypothalamic-pituitary-adrenal axes (HPA), while in behavioural patterns they display a proactive style, supporting the two-tier model (Koolhaas et al. 2010; Qu et al. 2018; Santicchia et al. 2022) and the labile relationships between glucocorticoid and behavioural variation (Duckworth et al. 2023). A meta-analysis across 21 species showed that individual hormone levels were only weakly correlated with proactive behavioural traits (aggression, boldness, exploration, and activity), explaining on average only 2% of the variation in personality (Niemelä, Dingemanse 2018). Moreover, a recent meta-analysis revealed that the direction of the link between stress regulation and behavioural traits depends on the species’ life history. In particular, in slow species, bolder individuals show lower stress reactivity, whereas in fast species, bolder individuals have higher stress reactivity (Duckworth et al. 2023). The midday gerbil is a small-sized fast-living species with a short life span, early maturation and intense reproduction (Tchabovsky et al. 2019a; Yang et al. 2020), which may explain why female colonists combine the potentiated HPA activity as an attribute of reactive coping style with a proactive bold behavioural strategy.

Links between dispersal propensity and behaviour were found in females but not in males also in great tits (Dingenmance et al. 2003), and females were, on average, more consistent in behavioural traits (excluding mate preferences) than males in personality studies (Bell et al. 2009). We do not exactly know whether the revealed behavioural patterns of female colonists are true personality traits – we had only a few repeated measures and, thus, could not partition within- and among-individual variation to assess individual consistency in behavioural responses (Brommer 2013; Dingemanse, Dochtermann 2013). However, previously, we have shown that, in the midday gerbils, behavioural traits (including boldness/exploration syndrome, sociability and trial execution speed) assessed in the same test procedure in the laboratory were highly repeatable and consistent across time and contexts, indicating personalities (Tchabovsky et al. 2019b; Tchabovsky et al. 2024). Combined with the strong consistency of behavioural profiles of female colonists across generations, this supports that the colonizer syndrome may be a heritable personality trait of females.

One more important inference from our study is that real-time observations allow recording short-term changes in the behaviour of colonists, which otherwise can be missed. If we had not observed the colonization process from the beginning when the first settlers arrived, we could have missed an abrupt change in male colonist behaviour after just one year post-colonies foundation and made wrong inferences about the sex-specific behavioural dynamics post-colonization. Missed short-term effects hinder detecting the flexibility of behavioural responses to changes in environmental uncertainty pre-, during, and post-colony foundation, calling for more studies conducted in real time, which can provide new insights into understanding the mechanisms of colonization.

## Supporting information

S1 File. Dataset for primary measurements

S2 File Dataset for composite variables

S3 File. Codes

## Supplementary Materials

**S1 File.** Dataset for primary measurements (pers_field_primary.csv)

**S2 File** Dataset for composite variables (pers_field_ad_pca.csv)

**S3 File.** Codes (field_tests_codes.R).

## Ethics

All applicable international and national guidelines for the care and use of animals were followed. All procedures conform to the ASAB/ABS Guidelines for the Treatment of Animals in Behavioural Research and Teaching (ASAB/ABS 2024). The research protocols for this study were approved by the Animal Ethics Commission of Severtsov Institute of Ecology and Evolution (protocol #58-2022).

## Data Accessibility

The datasets generated and analysed during the current study and codes are available as electronic Supplementary Materials.

## Authors’ Contributions

A.V.T.: Conceptualization, Formal analysis, Funding acquisition, Investigation, Methodology, Project administration, Resources, Supervision, Validation, Visualization, Writing – original draft, Writing – review and editing.

E.N.S.: Data curation, Investigation, Methodology, Writing – review and editing.

L.E.S.: Investigation, Methodology, Writing – review and editing.

I.S.K.: Data curation, Investigation, Writing – review and editing.

All authors gave final approval for publication and agreed to be held accountable for the work performed therein.

## Acknowledgments

We thank N. Ovchinnikova, D. Pozhariskii, Y. Chabovskaya, D.B. Vasiliev, V. Shved, Y. Vasilieva, A. Raspopova, A. Mokrousov, T. Demidova, A.B. Vasilieva, V. Trunov, A. Kalinin, V. Brukhno, A. Bogatchuk for assistance in field studies.

