## Supplementary material for "Flexible males, proactive females: personality tests show increased boldness/exploration in colonists damping with time in males but not in females": S3 File. Codes: field_tests_codes.R

```
#LME for primary measurements: source versus new colonies
#Field tests by gen_2
pers_field <- read.csv("D:/Work/R/data/pers_field_primary.csv",header=TRUE)
str(pers_field)


pers_field$animal_id<- as.factor(pers_field$animal_id)
pers_field$gen_2<- as.factor(pers_field$gen_2)


#MALES AD
#Subset: mad, spr, gen_2 == "s" OR gen_2 == "nc"
pers_field_mad <- subset(pers_field, sex_age == "mad" & season == "spr" & 
(gen_2 == "nc" | gen_2 == "s"))

str(pers_field_mad)

install.packages("meantables")
install.packages("MuMIn")

library(nlme)
library(emmeans)
library(car)

#IMMOB
#LME from nlme for immob from gen_2 and animal_id as RANDOM
pers_field_mad_immob <- lme(immob ~ gen_2, data = pers_field_mad, random = ~ 1 | animal_id, method="REML")
#str(pers_field_mad_immob)
summary(pers_field_mad_immob)


#GLS from nlme for the model with FIXED effects only
pers_field_mad_immob_fixed <- gls(immob ~ gen_2, data = pers_field_mad, method="REML")

#Comparing models WITH and WITHOUT (only fixed effects) random factor 
anova(pers_field_mad_immob, pers_field_mad_immob_fixed) 

#Means with 95%CI
pers_field_mad_immob_emm <-  emmeans(pers_field_mad_immob, ~ gen_2)
summary(pers_field_mad_immob_emm)


#OUT-2 EXCLUDED
#BO_ST
#LME from nlme for bo_st from gen_2 and animal_id as RANDOM

#Subset: mad, spr, note_st == "in" gen_2 == "s" OR gen_2 == "nc"
pers_field_mad <- subset(pers_field, sex_age == "mad" & season == "spr" & note_st == "in" & (gen_2 == "nc" | gen_2 == "s"))

#Selecting responce variable from the dataset
bo_st <- c("bo_st", "sex_age", "season", "note_st", "gen_2", "animal_id")
pers_field_mad_bo_st <- pers_field_mad [bo_st]
str(pers_field_mad_bo_st)

#Excluding Missing Values
pers_field_mad_bo_st_no_na <- na.omit(pers_field_mad_bo_st)
str(pers_field_mad_bo_st_no_na)

#LME with animal_id as Random factor
pers_field_mad_bo_st <- lme(bo_st ~ gen_2, data = pers_field_mad_bo_st_no_na, random = ~ 1 | animal_id, method="REML")
#str(pers_field_mad_bo_st)
summary(pers_field_mad_bo_st)


#GLS from nlme for the model with FIXED effects only
pers_field_mad_bo_st_fixed <- gls(bo_st ~ gen_2, data = pers_field_mad_bo_st_no_na, method="REML")

#Comparing models WITH and WITHOUT (only fixed effects)random factor 
anova(pers_field_mad_bo_st, pers_field_mad_bo_st_fixed) 

#Means with 95%CI
pers_field_mad_bo_st_emm <-  emmeans(pers_field_mad_bo_st, ~ gen_2)
summary(pers_field_mad_bo_st_emm)


#CO
#LME from nlme for co from gen_2 and animal_id as RANDOM

#Subset: mad, spr, note_st == "in", gen_2 == "s" OR gen_2 == "nc"
pers_field_mad <- subset(pers_field, sex_age == "mad" & season == "spr" & note_st == "in" & (gen_2 == "nc" | gen_2 == "s"))

#Selecting responce variable from the dataset
co <- c("co", "sex_age", "season", "note_st", "gen_2", "animal_id")
pers_field_mad_co <- pers_field_mad [co]
str(pers_field_mad_co)

#Excluding Missing Values
pers_field_mad_co_no_na <- na.omit(pers_field_mad_co)
str(pers_field_mad_co_no_na)

#LME with animal_id as Random factor
pers_field_mad_co <- lme(co ~ gen_2, data = pers_field_mad_co_no_na, random = ~ 1 | animal_id, method="REML")
#str(pers_field_mad_co)
summary(pers_field_mad_co)


#GLS from nlme for the model with FIXED effects only
pers_field_mad_co_fixed <- gls(co ~ gen_2, data = pers_field_mad_co_no_na, method="REML")

#Comparing models WITH and WITHOUT (only fixed effects)random factor 
anova(pers_field_mad_co, pers_field_mad_co_fixed) 

#Means with 95%CI
pers_field_mad_co_emm <-  emmeans(pers_field_mad_co, ~ gen_2)
summary(pers_field_mad_co_emm)


#BO_EP
#LME from nlme for bo_ep from gen_2 and animal_id as RANDOM

#Subset: mad, spr, note_ep == "in", gen_2 == "s" OR gen_2 == "nc"
pers_field_mad <- subset(pers_field, sex_age == "mad" & season == "spr" & note_ep == "in" & (gen_2 == "nc" | gen_2 == "s"))

#Selecting responce variable from the dataset
bo_ep <- c("bo_ep", "sex_age", "season", "note_ep", "gen_2", "animal_id")
pers_field_mad_bo_ep <- pers_field_mad [bo_ep]
str(pers_field_mad_bo_ep)

#Excluding Missing Values
pers_field_mad_bo_ep_no_na <- na.omit(pers_field_mad_bo_ep)
str(pers_field_mad_bo_ep_no_na)

#LME with animal_id as Random factor
pers_field_mad_bo_ep <- lme(bo_ep ~ gen_2, data = pers_field_mad_bo_ep_no_na, random = ~ 1 | animal_id, method="REML")
#str(pers_field_mad_bo_ep)
summary(pers_field_mad_bo_ep)


#GLS from nlme for the model with FIXED effects only
pers_field_mad_bo_ep_fixed <- gls(bo_ep ~ gen_2, data = pers_field_mad_bo_ep_no_na, method="REML")

#Comparing models WITH and WITHOUT (only fixed effects)random factor 
anova(pers_field_mad_bo_ep, pers_field_mad_bo_ep_fixed) 

#Means with 95%CI
pers_field_mad_bo_ep_emm <-  emmeans(pers_field_mad_bo_ep, ~ gen_2)
summary(pers_field_mad_bo_ep_emm)


#CD
#LME from nlme for cd from gen_2 and animal_id as RANDOM

#Subset: mad, spr, note_ep == "in", gen_2 == "s" OR gen_2 == "nc"
pers_field_mad <- subset(pers_field, sex_age == "mad" & season == "spr" & note_ep == "in" & (gen_2 == "nc" | gen_2 == "s"))

#Selecting responce variable from the dataset
cd <- c("cd", "sex_age", "season", "note_ep", "gen_2", "animal_id")
pers_field_mad_cd <- pers_field_mad [cd]
str(pers_field_mad_cd)

#Excluding Missing Values
pers_field_mad_cd_no_na <- na.omit(pers_field_mad_cd)
str(pers_field_mad_cd_no_na)

#LME with animal_id as Random factor
pers_field_mad_cd <- lme(cd ~ gen_2, data = pers_field_mad_cd_no_na, random = ~ 1 | animal_id, method="REML")
#str(pers_field_mad_cd)
summary(pers_field_mad_cd)


#GLS from nlme for the model with FIXED effects only
pers_field_mad_cd_fixed <- gls(cd ~ gen_2, data = pers_field_mad_cd_no_na, method="REML")

#Comparing models WITH and WITHOUT (only fixed effects)random factor 
anova(pers_field_mad_cd, pers_field_mad_cd_fixed) 

#Means with 95%CI
pers_field_mad_cd_emm <-  emmeans(pers_field_mad_cd, ~ gen_2)
summary(pers_field_mad_cd_emm)


#BO_STR
#LME from nlme for bo_str from gen_2 and animal_id as RANDOM

#Subset: mad, spr, note_str == "in", gen_2 == "s" OR gen_2 == "nc"
pers_field_mad <- subset(pers_field, sex_age == "mad" & season == "spr" & note_str == "in" & (gen_2 == "nc" | gen_2 == "s"))

#Selecting responce variable from the dataset
bo_str <- c("bo_str", "sex_age", "season", "note_str", "gen_2", "animal_id")
pers_field_mad_bo_str <- pers_field_mad [bo_str]
str(pers_field_mad_bo_str)

#Excluding Missing Values
pers_field_mad_bo_str_no_na <- na.omit(pers_field_mad_bo_str)
str(pers_field_mad_bo_str_no_na)

#LME with animal_id as Random factor
pers_field_mad_bo_str <- lme(bo_str ~ gen_2, data = pers_field_mad_bo_str_no_na, random = ~ 1 | animal_id, method="REML")
#str(pers_field_mad_bo_str)
summary(pers_field_mad_bo_str)


#GLS from nlme for the model with FIXED effects only
pers_field_mad_bo_str_fixed <- gls(bo_str ~ gen_2, data = pers_field_mad_bo_str_no_na, method="REML")

#Comparing models WITH and WITHOUT (only fixed effects)random factor 
anova(pers_field_mad_bo_str, pers_field_mad_bo_str_fixed) 

#Means with 95%CI
pers_field_mad_bo_str_emm <-  emmeans(pers_field_mad_bo_str, ~ gen_2)
summary(pers_field_mad_bo_str_emm)


#CS
#LME from nlme for cs from gen_2 and animal_id as RANDOM

#Subset: mad, spr, note_str == "in", gen_2 == "s" OR gen_2 == "nc"
pers_field_mad <- subset(pers_field, sex_age == "mad" & season == "spr" & note_str == "in" & (gen_2 == "nc" | gen_2 == "s"))

#Selecting responce variable from the dataset
cs <- c("cs", "sex_age", "season", "note_str", "gen_2", "animal_id")
pers_field_mad_cs <- pers_field_mad [cs]
str(pers_field_mad_cs)

#Excluding Missing Values
pers_field_mad_cs_no_na <- na.omit(pers_field_mad_cs)
str(pers_field_mad_cs_no_na)

#LME with animal_id as Random factor
pers_field_mad_cs <- lme(cs ~ gen_2, data = pers_field_mad_cs_no_na, random = ~ 1 | animal_id, method="REML")
#str(pers_field_mad_cs)
summary(pers_field_mad_cs)


#GLS from nlme for the model with FIXED effects only
pers_field_mad_cs_fixed <- gls(cs ~ gen_2, data = pers_field_mad_cs_no_na, method="REML")

#Comparing models WITH and WITHOUT (only fixed effects)random factor 
anova(pers_field_mad_cs, pers_field_mad_cs_fixed) 

#Means with 95%CI
pers_field_mad_cs_emm <-  emmeans(pers_field_mad_cs, ~ gen_2)
summary(pers_field_mad_cs_emm)


#TOT_DUR
#LME from nlme for cs from gen_2 and animal_id as RANDOM

#Subset: mad, spr, gen_2 == "s" OR gen_2 == "nc"
pers_field_mad <- subset(pers_field, sex_age == "mad" & season == "spr" & (gen_2 == "nc" | gen_2 == "s"))

#Selecting responce variable from the dataset
tot_dur <- c("tot_dur", "sex_age", "season", "gen_2", "animal_id")
pers_field_mad_tot_dur <- pers_field_mad [tot_dur]
str(pers_field_mad_tot_dur)

#Excluding Missing Values
pers_field_mad_tot_dur_no_na <- na.omit(pers_field_mad_tot_dur)
str(pers_field_mad_tot_dur_no_na)

#LME with animal_id as Random factor
pers_field_mad_tot_dur <- lme(tot_dur ~ gen_2, data = pers_field_mad_tot_dur_no_na, random = ~ 1 | animal_id, method="REML")
#str(pers_field_mad_tot_dur)
summary(pers_field_mad_tot_dur)


#GLS from nlme for the model with FIXED effects only
pers_field_mad_tot_dur_fixed <- gls(tot_dur ~ gen_2, data = pers_field_mad_tot_dur_no_na, method="REML")

#Comparing models WITH and WITHOUT (only fixed effects)random factor 
anova(pers_field_mad_tot_dur, pers_field_mad_tot_dur_fixed) 

#Means with 95%CI
pers_field_mad_tot_dur_emm <-  emmeans(pers_field_mad_tot_dur, ~ gen_2)
summary(pers_field_mad_tot_dur_emm)


#FEMALES AD
#OUT-2 EXCLUDED
pers_field <- read.csv("D:/Work/R/data/pers_field.csv",header=TRUE)

#Subset: fad, spr, gen_2 == "s" OR gen_2 == "nc"
pers_field_fad <- subset(pers_field, sex_age == "fad" & season == "spr" & (gen_2 == "nc" | gen_2 == "s"))

str(pers_field_fad)


#IMMOB
#LME from nlme for immob from gen_2 and animal_id as RANDOM
pers_field_fad_immob <- lme(immob ~ gen_2, data = pers_field_fad, random = ~ 1 | animal_id, method="REML")
str(pers_field_fad_immob)
summary(pers_field_fad_immob)


#GLS from nlme for the model with FIXED effects only
pers_field_fad_immob_fixed <- gls(immob ~ gen_2, data = pers_field_fad, method="REML")

#Comparing models WITH and WITHOUT (only fixed effects)random factor 
anova(pers_field_fad_immob, pers_field_fad_immob_fixed) 

#Means with 95%CI
pers_field_fad_immob_emm <-  emmeans(pers_field_fad_immob, ~ gen_2)
summary(pers_field_fad_immob_emm)


#BO_ST
#LME from nlme for bo_st from gen_2 and animal_id as RANDOM

#Subset: fad, spr, note_st == "in", gen_2 == "s" OR gen_2 == "nc"
pers_field_fad <- subset(pers_field, sex_age == "fad" & season == "spr" & note_st == "in" & (gen_2 == "nc" | gen_2 == "s"))

#Selecting responce variable from the dataset
bo_st <- c("bo_st", "sex_age", "season", "note_st", "gen_2", "animal_id")
pers_field_fad_bo_st <- pers_field_fad [bo_st]
str(pers_field_fad_bo_st)

#Excluding Missing Values
pers_field_fad_bo_st_no_na <- na.omit(pers_field_fad_bo_st)
str(pers_field_fad_bo_st_no_na)

#LME with animal_id as Random factor
pers_field_fad_bo_st <- lme(bo_st ~ gen_2, data = pers_field_fad_bo_st_no_na, random = ~ 1 | animal_id, method="REML")
#str(pers_field_fad_bo_st)
summary(pers_field_fad_bo_st)


#GLS from nlme for the model with FIXED effects only
pers_field_fad_bo_st_fixed <- gls(bo_st ~ gen_2, data = pers_field_fad_bo_st_no_na, method="REML")

#Comparing models WITH and WITHOUT (only fixed effects)random factor 
anova(pers_field_fad_bo_st, pers_field_fad_bo_st_fixed) 

#Means with 95%CI
pers_field_fad_bo_st_emm <-  emmeans(pers_field_fad_bo_st, ~ gen_2)
summary(pers_field_fad_bo_st_emm)


#CO
#LME from nlme for co from gen_2 and animal_id as RANDOM

#Subset: fad, spr, note_st == "in", gen_2 == "s" OR gen_2 == "nc"
pers_field_fad <- subset(pers_field, sex_age == "fad" & season == "spr" & note_st == "in" & (gen_2 == "nc" | gen_2 == "s"))

#Selecting responce variable from the dataset
co <- c("co", "sex_age", "season", "note_st", "gen_2", "animal_id")
pers_field_fad_co <- pers_field_fad [co]
str(pers_field_fad_co)

#Excluding Missing Values
pers_field_fad_co_no_na <- na.omit(pers_field_fad_co)
str(pers_field_fad_co_no_na)

#LME with animal_id as Random factor
pers_field_fad_co <- lme(co ~ gen_2, data = pers_field_fad_co_no_na, random = ~ 1 | animal_id, method="REML")
#str(pers_field_fad_co)
summary(pers_field_fad_co)


#GLS from nlme for the model with FIXED effects only
pers_field_fad_co_fixed <- gls(co ~ gen_2, data = pers_field_fad_co_no_na, method="REML")

#Comparing models WITH and WITHOUT (only fixed effects)random factor 
anova(pers_field_fad_co, pers_field_fad_co_fixed) 

#Means with 95%CI
pers_field_fad_co_emm <-  emmeans(pers_field_fad_co, ~ gen_2)
summary(pers_field_fad_co_emm)


#BO_EP
#LME from nlme for bo_ep from gen_2 and animal_id as RANDOM

#Subset: fad, spr, note_ep == "in", gen_2 == "s" OR gen_2 == "nc"
pers_field_fad <- subset(pers_field, sex_age == "fad" & season == "spr" & note_ep == "in" & (gen_2 == "nc" | gen_2 == "s"))

#Selecting responce variable from the dataset
bo_ep <- c("bo_ep", "sex_age", "season", "note_ep", "gen_2", "animal_id")
pers_field_fad_bo_ep <- pers_field_fad [bo_ep]
str(pers_field_fad_bo_ep)

#Excluding Missing Values
pers_field_fad_bo_ep_no_na <- na.omit(pers_field_fad_bo_ep)
str(pers_field_fad_bo_ep_no_na)

#LME with animal_id as Random factor
pers_field_fad_bo_ep <- lme(bo_ep ~ gen_2, data = pers_field_fad_bo_ep_no_na, random = ~ 1 | animal_id, method="REML")
#str(pers_field_fad_bo_ep)
summary(pers_field_fad_bo_ep)


#GLS from nlme for the model with FIXED effects only
pers_field_fad_bo_ep_fixed <- gls(bo_ep ~ gen_2, data = pers_field_fad_bo_ep_no_na, method="REML")

#Comparing models WITH and WITHOUT (only fixed effects)random factor 
anova(pers_field_fad_bo_ep, pers_field_fad_bo_ep_fixed) 

#Means with 95%CI
pers_field_fad_bo_ep_emm <-  emmeans(pers_field_fad_bo_ep, ~ gen_2)
summary(pers_field_fad_bo_ep_emm)


#CD
#LME from nlme for cd from gen_2 and animal_id as RANDOM

#Subset: fad, spr, note_ep == "in", gen_2 == "s" OR gen_2 == "nc"
pers_field_fad <- subset(pers_field, sex_age == "fad" & season == "spr" & note_ep == "in" & (gen_2 == "nc" | gen_2 == "s"))

#Selecting responce variable from the dataset
cd <- c("cd", "sex_age", "season", "note_ep", "gen_2", "animal_id")
pers_field_fad_cd <- pers_field_fad [cd]
str(pers_field_fad_cd)

#Excluding Missing Values
pers_field_fad_cd_no_na <- na.omit(pers_field_fad_cd)
str(pers_field_fad_cd_no_na)

#LME with animal_id as Random factor
pers_field_fad_cd <- lme(cd ~ gen_2, data = pers_field_fad_cd_no_na, random = ~ 1 | animal_id, method="REML")
#str(pers_field_fad_cd)
summary(pers_field_fad_cd)


#GLS from nlme for the model with FIXED effects only
pers_field_fad_cd_fixed <- gls(cd ~ gen_2, data = pers_field_fad_cd_no_na, method="REML")

#Comparing models WITH and WITHOUT (only fixed effects)random factor 
anova(pers_field_fad_cd, pers_field_fad_cd_fixed) 

#Means with 95%CI
pers_field_fad_cd_emm <-  emmeans(pers_field_fad_cd, ~ gen_2)
summary(pers_field_fad_cd_emm)


#BO_STR
#LME from nlme for bo_str from gen_2 and animal_id as RANDOM

#Subset: fad, spr, note_str == "in", gen_2 == "s" OR gen_2 == "nc"
pers_field_fad <- subset(pers_field, sex_age == "fad" & season == "spr" & note_str == "in" & (gen_2 == "nc" | gen_2 == "s"))

#Selecting responce variable from the dataset
bo_str <- c("bo_str", "sex_age", "season", "note_str", "gen_2", "animal_id")
pers_field_fad_bo_str <- pers_field_fad [bo_str]
str(pers_field_fad_bo_str)

#Excluding Missing Values
pers_field_fad_bo_str_no_na <- na.omit(pers_field_fad_bo_str)
str(pers_field_fad_bo_str_no_na)

#LME with animal_id as Random factor
pers_field_fad_bo_str <- lme(bo_str ~ gen_2, data = pers_field_fad_bo_str_no_na, random = ~ 1 | animal_id, method="REML")
#str(pers_field_fad_bo_str)
summary(pers_field_fad_bo_str)


#GLS from nlme for the model with FIXED effects only
pers_field_fad_bo_str_fixed <- gls(bo_str ~ gen_2, data = pers_field_fad_bo_str_no_na, method="REML")

#Comparing models WITH and WITHOUT (only fixed effects)random factor 
anova(pers_field_fad_bo_str, pers_field_fad_bo_str_fixed) 

#Means with 95%CI
pers_field_fad_bo_str_emm <-  emmeans(pers_field_fad_bo_str, ~ gen_2)
summary(pers_field_fad_bo_str_emm)


#CS
#LME from nlme for cs from gen_2 and animal_id as RANDOM

#Subset: fad, spr, note_str == "in", gen_2 == "s" OR gen_2 == "nc"
pers_field_fad <- subset(pers_field, sex_age == "fad" & season == "spr" & note_str == "in" & (gen_2 == "nc" | gen_2 == "s"))

#Selecting responce variable from the dataset
cs <- c("cs", "sex_age", "season", "note_str", "gen_2", "animal_id")
pers_field_fad_cs <- pers_field_fad [cs]
str(pers_field_fad_cs)

#Excluding Missing Values
pers_field_fad_cs_no_na <- na.omit(pers_field_fad_cs)
str(pers_field_fad_cs_no_na)

#LME with animal_id as Random factor
pers_field_fad_cs <- lme(cs ~ gen_2, data = pers_field_fad_cs_no_na, random = ~ 1 | animal_id, method="REML")
#str(pers_field_fad_cs)
summary(pers_field_fad_cs)


#GLS from nlme for the model with FIXED effects only
pers_field_fad_cs_fixed <- gls(cs ~ gen_2, data = pers_field_fad_cs_no_na, method="REML")

#Comparing models WITH and WITHOUT (only fixed effects)random factor 
anova(pers_field_fad_cs, pers_field_fad_cs_fixed) 

#Means with 95%CI
pers_field_fad_cs_emm <-  emmeans(pers_field_fad_cs, ~ gen_2)
summary(pers_field_fad_cs_emm)


#TOT_DUR
#LME from nlme for cs from gen_2 and animal_id as RANDOM

#Subset: fad, spr, gen_2 == "s" OR gen_2 == "nc"
pers_field_fad <- subset(pers_field, sex_age == "fad" & season == "spr" & (gen_2 == "nc" | gen_2 == "s"))

#Selecting responce variable from the dataset
tot_dur <- c("tot_dur", "sex_age", "season", "gen_2", "animal_id")
pers_field_fad_tot_dur <- pers_field_fad [tot_dur]
str(pers_field_fad_tot_dur)

#Excluding Missing Values
pers_field_fad_tot_dur_no_na <- na.omit(pers_field_fad_tot_dur)
str(pers_field_fad_tot_dur_no_na)

#LME with animal_id as Random factor
pers_field_fad_tot_dur <- lme(tot_dur ~ gen_2, data = pers_field_fad_tot_dur_no_na, random = ~ 1 | animal_id, method="REML")
#str(pers_field_fad_tot_dur)
summary(pers_field_fad_tot_dur)


#GLS from nlme for the model with FIXED effects only
pers_field_fad_tot_dur_fixed <- gls(tot_dur ~ gen_2, data = pers_field_fad_tot_dur_no_na, method="REML")

#Comparing models WITH and WITHOUT (only fixed effects)random factor 
anova(pers_field_fad_tot_dur, pers_field_fad_tot_dur_fixed) 

#Means with 95%CI
pers_field_fad_tot_dur_emm <-  emmeans(pers_field_fad_tot_dur, ~ gen_2)
summary(pers_field_fad_tot_dur_emm)


#LME for PCs by Generations
#Field tests PC by gen_2_cat
pers_field <- read.csv("D:/Work/R/data/pers_field_ad_pca.csv",header=TRUE)
str(pers_field)


pers_field$animal_id<- as.factor(pers_field$animal_id)
pers_field$gen_2_cat<- as.factor(pers_field$gen_2_cat)


install.packages("meantables")
install.packages("MuMIn")

library(nlme)
library(emmeans)
library(car)


#MALES AD -  PCs based on Separate PCA for Mad and Fad

#Subset: mad 
pers_field_mad <- subset(pers_field, sex_age == "mad")

#PC1_mad_fad
#LME from nlme for PC1_mad_fad from gen_2_cat and animal_id as RANDOM
pers_field_pc1_mad <- lme(pc_1_mad_fad ~ gen_2_cat, data = pers_field_mad, random = ~ 1 | animal_id, method="REML")
#str(pers_field_mad_pc1)
summary(pers_field_pc1_mad)


#Means with 95%CI
pers_field_pc1_mad_emm <-  emmeans(pers_field_pc1_mad, ~ gen_2_cat)
summary(pers_field_pc1_mad_emm)


#GLS from nlme for the model with FIXED effects only
pers_field_pc1_mad_fixed <- gls(pc_1_mad_fad ~ gen_2_cat, data = pers_field_mad, method="REML")

#Comparing models WITH and WITHOUT (only fixed effects)random factor 
anova(pers_field_pc1_mad, pers_field_pc1_mad_fixed) 


#PC2_mad_fad
#LME from nlme for PC2_mad_fad from gen_2_cat and animal_id as RANDOM
pers_field_pc2_mad <- lme(pc_2_mad_fad ~ gen_2_cat, data = pers_field_mad, random = ~ 1 | animal_id, method="REML")
#str(pers_field_mad_pc2)
summary(pers_field_pc2_mad)

#Means with 95%CI
pers_field_pc2_mad_emm <-  emmeans(pers_field_pc2_mad, ~ gen_2_cat)
summary(pers_field_pc2_mad_emm)


#GLS from nlme for the model with FIXED effects only
pers_field_pc2_mad_fixed <- gls(pc_2_mad_fad ~ gen_2_cat, data = pers_field_mad, method="REML")

#Comparing models WITH and WITHOUT (only fixed effects)random factor 
anova(pers_field_pc2_mad, pers_field_pc2_mad_fixed)


#PC3
#PC3_mad_fad
#LME from nlme for PC3_mad_fad from gen_2_cat and animal_id as RANDOM
pers_field_pc3_mad <- lme(pc_3_mad_fad ~ gen_2_cat, data = pers_field_mad, random = ~ 1 | animal_id, method="REML")
#str(pers_field_mad_pc3)
summary(pers_field_pc3_mad)


#Means with 95%CI
pers_field_pc3_mad_emm <-  emmeans(pers_field_pc3_mad, ~ gen_2_cat)
summary(pers_field_pc3_mad_emm)


#GLS from nlme for the model with FIXED effects only
pers_field_pc3_mad_fixed <- gls(pc_3_mad_fad ~ gen_2_cat, data = pers_field_mad, method="REML")

#Comparing models WITH and WITHOUT (only fixed effects)random factor 
anova(pers_field_pc3_mad, pers_field_pc3_mad_fixed) 
 


#FEMALES AD -  PCs based on Separate PCA for Mad and Fad

#Subset: fad 
pers_field_fad <- subset(pers_field, sex_age == "fad")

#PC1_mad_fad
#LME from nlme for PC1_mad_fad from gen_2_cat and animal_id as RANDOM
pers_field_pc1_fad <- lme(pc_1_mad_fad ~ gen_2_cat, data = pers_field_fad, random = ~ 1 | animal_id, method="REML")
#str(pers_field_pc1_fad)
summary(pers_field_pc1_fad)


#Means with 95%CI
pers_field_pc1_fad_emm <-  emmeans(pers_field_pc1_fad, ~ gen_2_cat)
summary(pers_field_pc1_fad_emm)


#GLS from nlme for the model with FIXED effects only
pers_field_pc1_fad_fixed <- gls(pc_1_mad_fad ~ gen_2_cat, data = pers_field_fad, method="REML")

#Comparing models WITH and WITHOUT (only fixed effects)random factor 
anova(pers_field_pc1_fad, pers_field_pc1_fad_fixed) 


#PC2_mad_fad
#LME from nlme for PC2_mad_fad from gen_2_cat and animal_id as RANDOM
pers_field_pc2_fad <- lme(pc_2_mad_fad ~ gen_2_cat, data = pers_field_fad, random = ~ 1 | animal_id, method="REML")
#str(pers_field_pc2_fad)
summary(pers_field_pc2_fad)


#Means with 95%CI
pers_field_pc2_fad_emm <-  emmeans(pers_field_pc2_fad, ~ gen_2_cat)
summary(pers_field_pc2_fad_emm)


#GLS from nlme for the model with FIXED effects only
pers_field_pc2_fad_fixed <- gls(pc_2_mad_fad ~ gen_2_cat, data = pers_field_fad, method="REML")

#Comparing models WITH and WITHOUT (only fixed effects)random factor 
anova(pers_field_pc2_fad, pers_field_pc2_fad_fixed) 


#PC3_mad_fad
#LME from nlme for PC3_mad_fad from gen_2_cat and animal_id as RANDOM
pers_field_pc3_fad <- lme(pc_3_mad_fad ~ gen_2_cat, data = pers_field_fad, random = ~ 1 | animal_id, method="REML")
#str(pers_field_pc3_fad)
summary(pers_field_pc3_fad)

#Means with 95%CI
pers_field_pc3_fad_emm <-  emmeans(pers_field_pc3_fad, ~ gen_2_cat)
summary(pers_field_pc3_fad_emm)


#GLS from nlme for the model with FIXED effects only
pers_field_pc3_fad_fixed <- gls(pc_3_mad_fad ~ gen_2_cat, data = pers_field_fad, method="REML")

#Comparing models WITH and WITHOUT (only fixed effects)random factor 
anova(pers_field_pc3_fad, pers_field_pc3_fad_fixed) 
```
